# P4HA2: A link between tumor-intrinsic hypoxia, partial EMT and collective migration

**DOI:** 10.1101/2022.02.28.482409

**Authors:** Vaishali Aggarwal, Sarthak Sahoo, Vera S. Donnenberg, Priyanka Chakraborty, Mohit Kumar Jolly, Shilpa Sant

## Abstract

Epithelial-to-mesenchymal transition (EMT), a well-established phenomenon studied across pan-cancer types, has long been known to be a major player in driving tumor invasion and metastasis. Recent studies have highlighted the importance of partial EMT phenotypes in metastasis [1]. Initially thought as a transitional state between epithelial and mesenchymal phenotypic states, partial EMT state is now widely recognized as a key driver of intra-tumoral heterogeneity and phenotypic plasticity, further accelerating tumor metastasis and therapeutic resistance. However, how tumor microenvironments regulate partial EMT phenotypes remains unclear. We have developed unique size-controlled three-dimensional microtumor models that recapitulate tumor-intrinsic hypoxia and the emergence of collectively migrating cells. In this study, we further interrogate these microtumor models to understand how tumor-intrinsic hypoxia regulates partial EMT and collective migration in hypoxic large microtumors fabricated from T47D breast cancer cells. We also compared global gene expression profiles of hypoxic, migratory microtumors to that of non-hypoxic, non-migratory microtumors at early and late time-points. Using our microtumor models, we identified unique gene signatures for tumor-intrinsic hypoxia (early *versus* late), partial EMT and migration (pre-migratory *versus* migratory phenotype). Through differential gene expression analysis between the microtumor models with an overlap of hypoxia, partial EMT and migration signatures, we identified prolyl 4-hydroxylase subunit 2 (P4HA2), a hypoxia responsive gene, as a central regulator common to hypoxia, partial EMT and collective migration. Further, the inhibition of P4HA2 significantly blocked collective migration in hypoxic microtumors. Thus, using the integrated computational-experimental analysis, we identify the key role of P4HA2 in tumor-intrinsic hypoxia-driven partial EMT and collective migration.

## 1. Introduction

Epithelial-to-mesenchymal transition (EMT) during which epithelial cells (E) lose their cell-cell adhesion properties and transition to a mesenchymal state (M), has long been known to drive tumor invasion and metastasis [2–4]. Historically, EMT was thought to have terminal E and M states and earlier studies focused on underlying mechanisms driving EMT. Emerging evidence now suggests the existence of partial EMT phenotype, a transitional state between the E and M states, wherein cells retain the epithelial phenotype with gain of mesenchymal phenotype [2, 5]. These E, M and partial EMT subsets of malignant cells are present concurrently during tumor progression as the cells transition from E to partial EMT and to M phenotype driving the intra-tumoral heterogeneity [6, 7]. However, molecular mechanisms that drive partial EMT states, and thereby, intra-tumoral heterogeneity, tumor progression, and therapy resistance remain understudied.

It is widely accepted that the tumor microenvironment (TME) plays a critical role in tumor progression [8]. TME is shaped by tumor-intrinsic factors (*e.g*., hypoxia, acidic microenvironment) and tumor-extrinsic factors (extracellular matrix and stromal related factors) [8]. TME consists of heterogeneous cellular composition including malignant cells, endothelial cells, stromal cells, fibroblasts, immune cells and extracellular matrix [8, 9]. Of these, tumor-intrinsic hypoxia has been shown to drive and stabilize partial EMT [6, 8, 10]. In oesophageal squamous cell carcinoma, malignant cells overexpressed hypoxia inducible factor 1 alpha (HIF1a) and exhibited partial EMT phenotype characterized by co-expression of E-cadherin (ECAD) and vimentin (VIM) [11]. Acidic microenvironment also contributed to partial EMT in MCF-7 cells, which co-expressed epithelial (ZO-1 and E-CAD) and mesenchymal (VIM, SNAI1 and β-catenin) markers [12]. In addition to tumor-intrinsic factors, tumor-extrinsic factors such as extracellular matrix (fibrinogen or collagen matrix) [13, 14], cancer-associated fibroblasts and stromal cells [15, 16] and tumor associated macrophages [17, 18] have all shown to drive and sustain partial EMT phenotype. These studies support the collective role of TME in inducing partial EMT tumor states.

To investigate mechanisms by which tumor-intrinsic hypoxia drives partial EMT phenotype, it is imperative to have experimental model systems, which can successfully recapitulate inherent tumor-intrinsic hypoxic microenvironment. Majority of published studies utilize exposure of 2D cell monolayers or three-dimensional (3D) spheroids to hypoxic chambers, where all cells in 2D monolayers or even outer cells in 3D spheroids are exposed to constant hypoxia, in contrast to the dynamic, spatial oxygen gradients that are observed *in vivo* [8]. Our lab has engineered a unique 3D microtumour models, which recapitulate tumor-intrinsic hypoxic microenvironment, solely driven by the size of microtumors without any genetic manipulation or any artificial culture conditions [10, 19–21]. A major advantage of microtumor models is the ease and reproducibility of generation of both, control, non-hypoxic, non-migratory microtumors (150μm) as well as hypoxic, migratory microtumors (600μm) *in vitro*. We have shown that migratory phenotype is easily induced in large hypoxic microtumors made of parent non-migratory T47D cells without any external stimulus [19–21]. We have also observed the presence of heterogenous tumor states *i.e*., E, M and E/M (partial EMT) [21]. Specifically, we have shown that large hypoxic microtumors gain the expression of mesenchymal marker (VIM) between day 3 to 6 while maintaining epithelial marker (E-CAD) expression, supporting the acquisition of partial EMT phenotype [21]. Subsequent studies highlighted the co-localization of E-CAD+ VIM+ or CD90+CD44+CTK+ (partial EMT) tumor cells at the leading edge of the migratory front, further solidifying the role of partial EMT in driving collective migration [21, 22].

The current study delineates how tumor-intrinsic hypoxia drives partial EMT and collective migration. To investigate the potential mechanisms underlying tumor-intrinsic hypoxia-driven partial EMT phenotype, we comparted global gene expression profiles obtained from microarray platform of large hypoxic microtumors at early (day 1) and late time-points (day 6) with that of non-hypoxic small microtumors. We identified enrichment of hypoxia, partial EMT and migration gene signatures early on day 1 (600/D1) and at later time point (600/D6) in large microtumours. We identified *P4HA2*, a hypoxia-responsive gene as central regulator that showed overlap between hypoxia, partial EMT and migration gene signatures. Further the P4HA inhibition studies significantly abrogated migration in large microtumors supporting its role in partial EMT and collective migration.

## 2. Materials and methods

### 2.1 Cell culture

T47D cell line was purchased from American type culture collection (ATCC). Characterization and authentication of T47D cells was done at the University of Arizona Genomic Core facility using PowerPlex16HS PCR kit as previously described [20, 21]. All cell culture supplies and media were procured from Corning^®^ and Mediatech^®^, respectively, unless otherwise specified. T47D cells were passaged and maintained in T75 flask in Dulbecco’s modified Eagle medium (DMEM) complete growth media (MT10013CV, Corning^®^, USA) supplemented with 10% fetal bovine serum (FBS) (S11250, Atlanta Biologicals, USA) and 1% penicillin-streptomycin (300-002-CI, Corning^®^, USA) in a humidified incubator with 5% CO_2_ at 37°C. Cells were maintained at 40-60% confluency before seeding into size-controlled hydrogel microwell devices.

### 2.2 Generation of size-controlled microtumors

The size-controlled T47D microtumors with diameters 100-150 μm (‘small’ microtumors) and 450-600 μm (‘large’ microtumors) were microfabricated using non-adhesive polyethylene glycol (PEG) hydrogel microwell devices as previously described [10, 19–21]. Briefly, uniform size 150 μm and 600 μm hydrogel microwell devices (1×1 cm^2^) were fabricated using polydimethyl siloxane (PDMS) stamps containing posts with 1:1 aspect ratio of height: diameter. Polyethylene glycol dimethacrylate (PEGDMA, 1000Da, Polysciences Inc., USA) solution (20% w/v) with 1% w/v photoinitiator (Irgacure-1959, Ciba AG CH-4002, Switzerland) was photo-crosslinked under the PDMS stamps to from size-controlled hydrogel microwells using OmniCure S2000 curing station (200W Lamp, 5W/cm^2^, EXFO, Canada). The fabricated hydrogel microwell devices were sterilized in 70% ethanol under UV light in laminar hood for 1h. Post-sterilization, devices were washed thrice with Dulbecco’s phosphate buffered saline (DPBS) without calcium and magnesium (#21-031-CV, Corning™, USA). Subsequently, T47D cells (1.0×10^6^ T47D cells/50μL/device) were seeded on to 1×1 cm^2^ hydrogel microwell devices. Excess cells remaining on top of the gels were washed slowly with DPBS. These cell-seeded microwell devices were then cultured in a humidified incubator with 5% CO_2_ at 37°C for six days with replacement of 50% media every day.

To study how tumor-intrinsic hypoxia drives partial EMT phenotype, large 600μm microtumors were cultured over 6-day period. The small non-hypoxic and non-migratory 150μm microtumors [20, 21] served as controls.

### 2.3 Multicolor flow cytometry staining

Small and large microtumors were cultured as described under section 2.2. At end of day 1, 3, and 6 of the culture, the microtumors were collected, washed with DBPS and disaggregated by incubating in Trypsin-EDTA for 5-10 minutes. For flow cytometry, three devices of 150μm microtumors and 1 device of 600μm microtumors were pooled to obtain required number of cells for each replicate and 3 such biological replicates for each group (150/D1, 150/D3, 150/D6, 600/D1, 600/D3, 600/D6) were used. The disaggregated cells were resuspended in PBS and processed for flow cytometric staining. Non-specific binding of fluorochrome-conjugated antibodies was minimized by pre-incubating pelleted cell suspensions for 5 min with neat decomplemented (56°C, 30 min) mouse serum (5 μL) [23]. Prior to intracellular cytokeratin or intracellular E-Cadherin staining, cells were stained for surface E-Cadherin (Biolegend, Cat. No. 147315) and fixed with 2% methanol-free formaldehyde (Polysciences, Warrington, PA). Cells were then permeabilized with 0.1% saponin (Beckman Coulter) in DPBS with 0.5% human serum albumin (10 min at room temperature). The permeabilized cell pellets were incubated with 5 μL of neat mouse serum for 5 min, centrifuged and decanted. The cell pellet was disaggregated and incubated with 2 μL of anti-pan cytokeratin-FITC (Beckman Coulter, Cat. No. IM2356U, Clone J1B3). A separate tube containing cells unstained for surface E-Cadherin was incubated after permeabilization with E-Cadherin antibody- (R&D Cat. No. FAB18381P, clone DECMA-1, ECD fluorochrome) to determine total (surface and intracellular) E-Cadherin expression. Finally, cells were resuspended to a concentration of 5 × 10^6^ cells/mL and DAPI (Invitrogen Cat. No. D1306) was added to the cells at a concentration of 10 μg/mL. Cells were acquired on Fortessa SORP flow cytometer (BD Biosciences, San Diego, CA). The instrument was calibrated using CS and T beads (BD Biosciences, Cat. No. 650621), and PMT voltages were adjusted to predetermined target channels using the seventh peak of 8-peak Rainbow Calibration Particles (Spherotech, Lake Forest, IL, Cat. No. RCP-30-5A;) as a reference point. FITC (BD Biosciences, Cat. No. 349502 for cytokeratin expression) and single stained BD CompBeads anti-mouse IGκ (BD Biosciences Cat. No. 51-90-9001229, for E-Cadherin-CF594 clone DECMA1 expression), and unstained cells (DAPI only) were used as spectral compensation standards [24].

#### Analysis of FCS files

For each sample, data were exported into FCS files and organized by experiment into playlists using VenturiOne software (Applied Cytometry Systems, Dinnington, UK, V7.3). The gate for marker positive events was determined such that all negative controls (isotypes) represented less than 1% of clean events. Analytical data (event count, percent positive, MFI) were exported to CSV files and imported into Excel, where they were associated with sample ID and reagent.

### 2.4 Treatment of microtumors with prolyl 4-hydroxylase (P4HA) inhibitor

To investigate the role of hypoxia-induced partial EMT on collective migration in large microtumors, we studied the effect of *P4HA2* inhibition, a common hypoxia and partial EMT marker identified through our bioinformatic analysis. *P4HA2* was inhibited by using 20 μmol/L 1,4-dihydrophenonthrolin-4-one-3-carboxylic acid (1,4-DPCA) (SC-200758, Santa Cruz, USA). 1,4-DPCA, is a selective inhibitor of prolyl 4-hydroxylase, which potentially targets HIF-1α upon hydroxylation of proline residue. Large 600μm microtumors were treated with 1,4-DPCA (20μM) from day 1 over six-day period with replenishment with every change of media. To evaluate the effect of *P4HA* inhibition (*P4HA2i*) on collective migration of microtumors, number of migrating and non-migrating microtumors were counted on day 3 and day 6 (control *versus* treated microtumor devices). In addition, the control *versus* treated microtumors were also imaged at 4x and 10x magnification (Zeiss Primo-vert) on day 3 and day 6, and distance of migration (d) was calculated as described previously [21] for evaluating migration kinetics. Post-treatment, microtumors were collected at day 3 and day 6, respectively from control (untreated) and *P4HA2i*-treated devices for mRNA isolation. [21, 25].

### 2.5 Quantitative real time-polymerase chain reaction (qRT-PCR)

To validate the data obtained from microarray experiment, we validated the expression changes using qRT-PCR (QuantStudio 3, Applied Biosystems, USA). For each experiment, microtumors cultured in 1X1 cm^2^ devices were harvested from PEG hydrogel devices (at least two devices for 600μm microtumors and four devices for 150μm microtumors) and stored in −80°C in Trizol. RNA was isolated using GeneJET RNA purification kit (#K0732, Thermo Scientific, USA) as per manufacturer’s protocol. DNase I (#EN0521, ThermoScientific, USA) treatment was used to remove genomic DNA from the RNA samples as per manufacturer’s protocol in thermocycler (T100™, Bio-Rad Laboratories, Inc., USA). RNA concentration was quantitated using Nanodrop (#840274200™, Thermo Scientific, USA) and RNA quality was determined by absorbance ratio at 260/280 nm. Primers for *GAPDH* (housekeeping gene), *P4HA1* and *P4HA2*, were procured from KiCqStart™ primers (H_GAPDH3_3, H_P4HA2_1 and H_P4HA2_2, Sigma-Aldrich, USA). mRNA expression levels of above-mentioned genes were estimated using iTaq™ Universal SYBR^®^ Green One-step qRT-PCR kit (#1725151, Bio-Rad Laboratories Inc., USA) per manufacturer’s protocol. Each reaction was run in triplicates (n=3) with no template control (NTC) and no reverse-transcriptase control (RTC). The complete experiment was repeated in triplicates (n=3). Relative fold change expression for individual genes was calculated using 2^−ΔΔCt^ method [26] and analyzed for fold change expression with respective 600/D6 untreated samples as controls.

### 2.6 Bioinformatic analysis of gene expression profiling data

To investigate the role of tumor-intrinsic hypoxia in driving phenotypic plasticity and collective migration, we have performed microarray analysis of 150/D6, 600/D1 and 600/D6 microtumors. [27]. In the current study, we analyzed microarray data deposited in the Gene Expression Omnibus database at the National Center for Biotechnology Information (**GSE166211**) [27] to establish the link between hypoxia, partial EMT and tissue migration by comparing 600/D1 versus 150/D6 and 600/D6 versus 150/D6 comparison groups.

#### Identification of differentially expressed genes

Microarray data deposited in the Gene Expression Omnibus database at the National Center for Biotechnology Information (GSE166211) [27] was used for the analysis reported in this study. The microarray analysis for identification of differentially expressed genes in 600/D1 *versus* 150/D6 as well as 600/D6 *versus* 150D6 was carried out as described previously [27]. The statistical analysis on each comparison was performed using Transcriptome Analysis Console (TAC 4.0, ThermoFisher Scientific, USA) using one-way ANOVA. A cut-off *p*-value of 0.01 was used for hierarchical clustering. Differentially expressed genes were considered significant using *p-value* ≤ 0.05, false discovery rate (FDR) ≤ 0.05, and fold change ± 2.

#### Statistical enrichment of hypoxia, partial EMT and tissue migration signatures

The gene ontology enrichment of hallmark hypoxia in 600/D1 *versus* 150/D6 and 600/D6 *versus* 150/D6 groups was carried out using the pipeline as described in our previous work [27]. We used partial EMT gene signatures previously identified for head and neck cancer [28] as a reference gene signature set to assess the partial EMT signatures in 600/D1 and 600/D6 microtumors, Gene ontology database for GO: Tissue migration (GO:0090130) [29, 30] was used to assess the tissue migratory signatures of the microtumors.

#### ssGSEA scoring of biological pathways

Hallmark signatures from MsigDB were used to quantify the activity of different biological pathways using the ssGSEA for scoring and quantification of the same [31]. The ssGSEA function [32] in the python package gseapy (https://github.com/zqfang/gseapy) was used for scoring of the biological pathways.

### 2.7 Statistical Analysis

Statistical analysis was done using GraphPad Prism (v9.0). Descriptive statistics and statistical comparisons (Student’s t-test, 2-tailed) were performed, and data are represented as mean ± SEM (standard error of mean) from three replicates. Unpaired *t*-test was used to compare two groups and *p*-value less than 0.05 was considered significant. P-values were Bonferroni-corrected for multiple comparisons.

## 3. Results

### 3.1 Tumor-intrinsic hypoxia is enriched in large microtumors compared to small microtumors

To understand the transcriptomic differences between 150/D6, 600/D1 and 600/D6 microtumors, we performed Principal Component Analysis (PCA) on the gene expression matrices containing three biological replicates for each of the three microtumor models. Upon plotting the samples on a two-dimensional PCA space, we found that ~56% of the variance was explained by the first two PCA components (**Figure 1A**). Furthermore, we found that the replicates clustered together on the PCA space with the inter-biological sample distances larger than the distances between replicates for a given biological sample. While 600/D6 microtumors varied from 150/D6 microtumors along the PCA-1 axis, the 600/D1 varied along both PCA-1 and PCA-2 with respect to either 150/D6 or 600/D6 samples. This difference indicates that some biological processes vary specifically in the 600/D1 and not in the 600/D6 samples, with 150/D6 samples as a control group. To investigate the biological processes that contribute to the variances in these biological samples, we calculated the ssGSEA scores for hallmark pathways for all the three microtumor groups. Upon doing a differential analysis of scores between the pairs of microtumor groups 600/D1 versus 150/D6 and 600/D6 versus 150/D6 (**Figure 1B**), we found that Hallmark Hypoxia was consistently upregulated in both these comparison sets, along with TNFa signaling, EMT, p53 signaling and unfolded protein response, although to varying degrees. This mapping shows that tumor-intrinsic hypoxia is upregulated as early as day 1 (600/D1 microtumors) and may contribute to initiate the migratory phenotype in these microtumors. To assess the extent of enrichment of hypoxia signature, we calculated the ssGSEA scores for the Hallmark Hypoxia and found that the hypoxia signature was significantly higher in the 600/D1 and 600/D6 compared to 150/D6 microtumors (**Figure 1C**).

**Figure 1:**
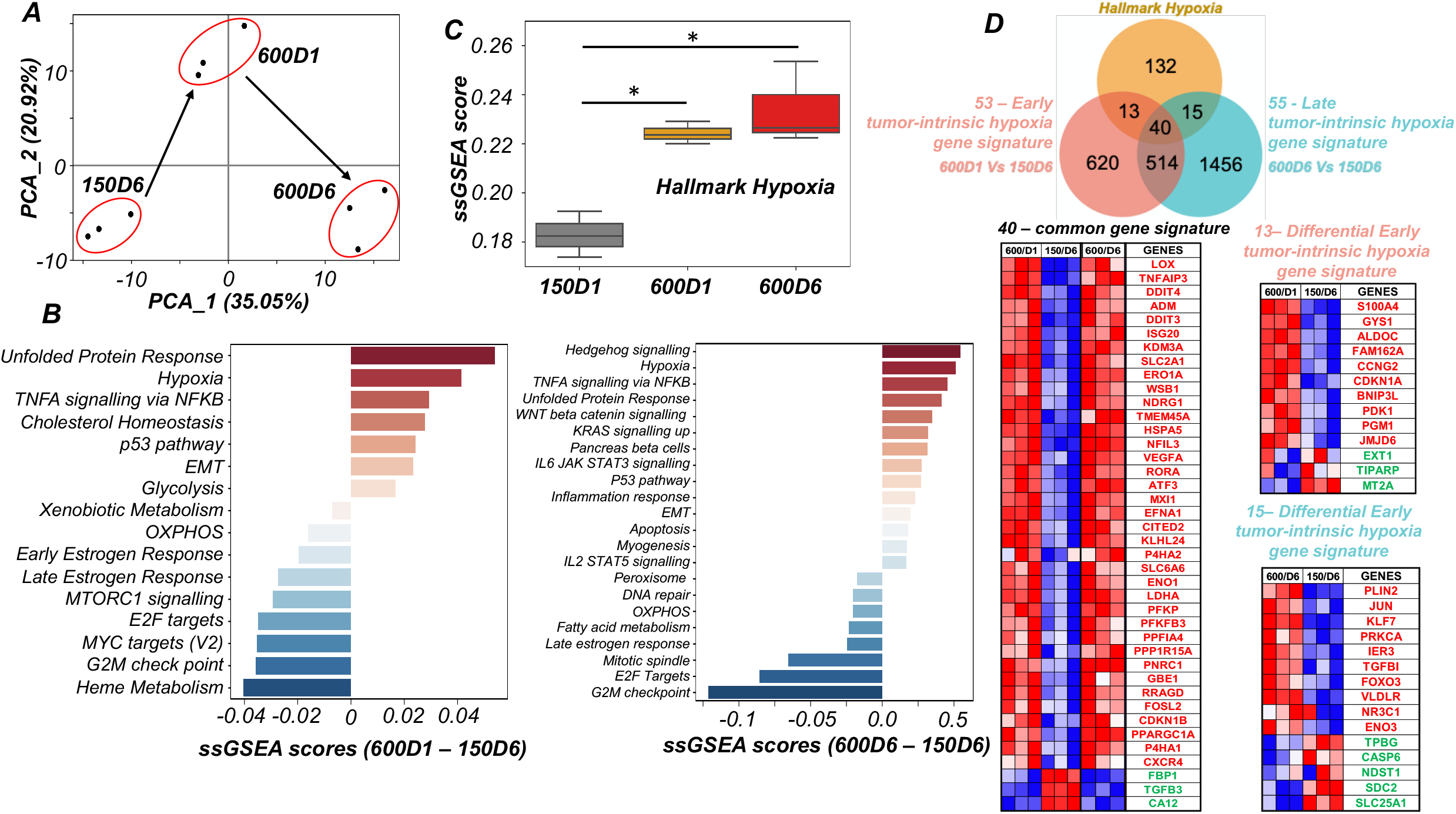
Tumor-intrinsic hypoxia is highly enriched in large microtumors. **(A)** Principal component analysis (PCA) plot for similarity assessment of gene expression levels across biological replicates of *150/D6, 600/D1* and *600/D6* microtumors fabricated from T47D cells; **(B)** ssGSEA enrichment of key biological processes in *600/D1 versus 150D6* and *600D6 versus 150D6* comparison groups. *Hallmark Hypoxia* is enriched as early as day 1 and maintained over six-day period. Only hallmark gene sets which show significant differences have been shown (*p*-value < 0.05; Student’s T-test); **(C)** ssGSEA score analysis for *Hallmark Hypoxia* in 600/D1 and 600/D6 groups individually with 150/D6 microtumors as control shows significant enrichment of tumor-intrinsic hypoxia in both 600/D1 and 600/D6 groups. **(D)** Venn diagram comparison of *Hallmark Hypoxia* with *600/D1 versus 150D6* and *600D6 versus 150D6* comparison groups shows an overlap of 53 genes for *600/D1* (denoted as *early tumor-intrinsic hypoxia gene signature*) and 55 genes for *600/D6 (denoted as late tumor-intrinsic hypoxia gene signature*). Forty genes are common among *Hallmark Hypoxia*, early and late tumor-intrinsic hypoxia gene signatures. Thirteen genes are differentially expressed only in *600/D1* and 15 genes are differentially expressed only in *600/D6*. These are represented as heat maps with corresponding intensity values of genes from biological replicates in 600/D1 and 600/D6 *versus* 150/D6 group. In the heatmaps, ‘*red*’ shows upregulated status and *‘blue’* shows downregulated status. The corresponding gene names are illustrated in *‘red’* for upregulated status and *‘green’* for downregulated status.

To obtain a comprehensive list of hypoxia gene signatures for large hypoxic microtumors (600/D1 and 600/D6) compared to small non-hypoxic microtumors (150/D6), we evaluated the overlap between Hallmark Hypoxia in 600/D1 *versus* 150/D6 and 600/D6 *versus* 150/D6 comparison groups (**Fig 1D**). The gene signatures from Hallmark Hypoxia were collated to generate comprehensive hypoxia gene signature sets for 600/D1 *versus* 150/D6 (hereafter referred as “early tumor-intrinsic hypoxia gene signature”) and 600/D6 *versus* 150/D6 (hereafter referred to as “late tumor-intrinsic hypoxia gene signature’). A total of 53 genes were enriched in 600/D1 versus 150/D6 resulting in early tumor-intrinsic hypoxia gene signature (**Figure 1D)** while late tumor-intrinsic hypoxia gene signature was enriched with 55 genes in 600/D6 versus 150/D6. The Venn diagram-based comparative analysis identified 13 genes differentially expressed only in 600/D1 and 15 genes expressed only in 600/D6 (**Figure 1D**). We also found 40 genes common to both ‘early’ and ‘late’ tumor-intrinsic hypoxia gene signatures that are differentially expressed as early as day 1. Of these 40 common genes, 37 genes were upregulated, and three genes were downregulated. Of the 13 genes differentially expressed only in 600/D1 (**Figure 1D**), *S100A4, GYS1, ALDOC, FAM162A* and *CCNG2* were most upregulated while *MT2A, TIRAP* and *EXT1* were top downregulated genes (in descending order of expression). Of these hypoxia gene signatures, elevated expression of *S100A4* has been correlated to hypoxia-induced tumor invasion and metastasis in esophageal squamous cell carcinoma and ovarian cancer [33, 34]. Of note are the hypoxia-induced genes *GYS1* [35, 36], *FAM162A* [37], *CCNG2* [38], *MT2A* [39, 40], which are induced due to hypoxic response across different cancer types; however, their role as hypoxia responsive genes in breast cancer is not yet reported. Note that in these studies, hypoxia was induced either using hypoxic chambers or with overexpression of HIF-1α, thus may not mimic naturally induced tumor-intrinsic hypoxia seen in our large microtumor models. In the ‘late tumor-intrinsic hypoxia’ signature of 600/D6 microtumors, *PLIN2* was the most upregulated gene followed by *JUN, KLF7, PRKCA* and *IER3* while *SLC25A1, SDC2, NDST1, CASP6* and *TPBG* were top five downregulated genes (in descending order of expression) (**Figure 1D**). Of these gene signatures, *PLIN2, JUN* and *PRKCA* are shown to be induced in response to hypoxia [41–43]. Taken together; gene expression analysis of large microtumor models (600/D1, 600/D6) with tumor size-induced natural hypoxia at day 1 and day 6 compared to non-hypoxic 150/D6 microtumors contributed ‘early’ and ‘late’ tumor-intrinsic hypoxia gene signatures.

### 3.2 Large microtumors exhibit enrichment of partial EMT signature in comparison to small microtumors

Our earlier work has demonstrated that the large 600μm microtumors with tumor-intrinsic hypoxia exhibit partial EMT phenotype characterized by gain of mesenchymal marker (VIM) without loss of epithelial marker (E-CAD) [21]. Based on these initial findings, we further investigated partial EMT signatures enriched by tumor-intrinsic hypoxia in large 600μm microtumors on day 1 and day 6.

First, we assessed changes in the epithelial and mesenchymal characteristics of 600/D1 and 600/D6 microtumors compared to control 150/D6 tumors. Specifically, we calculated the ssGSEA scores of epithelial genes as reported by Tan *et al*. 2014 [44] and observed that there was a robust decline in the epithelial nature in 600/D1 and 600/D6 compared to 150/D6 tumors (**Figure 2A**). Using the mesenchymal signatures from the same study [44], we found that the mesenchymal signature was upregulated significantly in 600/D6, but not in 600/D1 microtumors (**Figure 2B**). This could indicate that the 600/D1 tumors are more likely to be in a partial EMT phenotype and the 600/D6 tumors are more likely to be closer to a complete EMT phenotype. To validate this, we plotted the samples on a two-dimensional plane of epithelial and mesenchymal phenotype scores. Indeed, 600/D1 tumors lie in an intermediate region between the more epithelial 150/D6 and the more mesenchymal 600/D6 tumors (**Figure 2C**).

**Figure 2:**
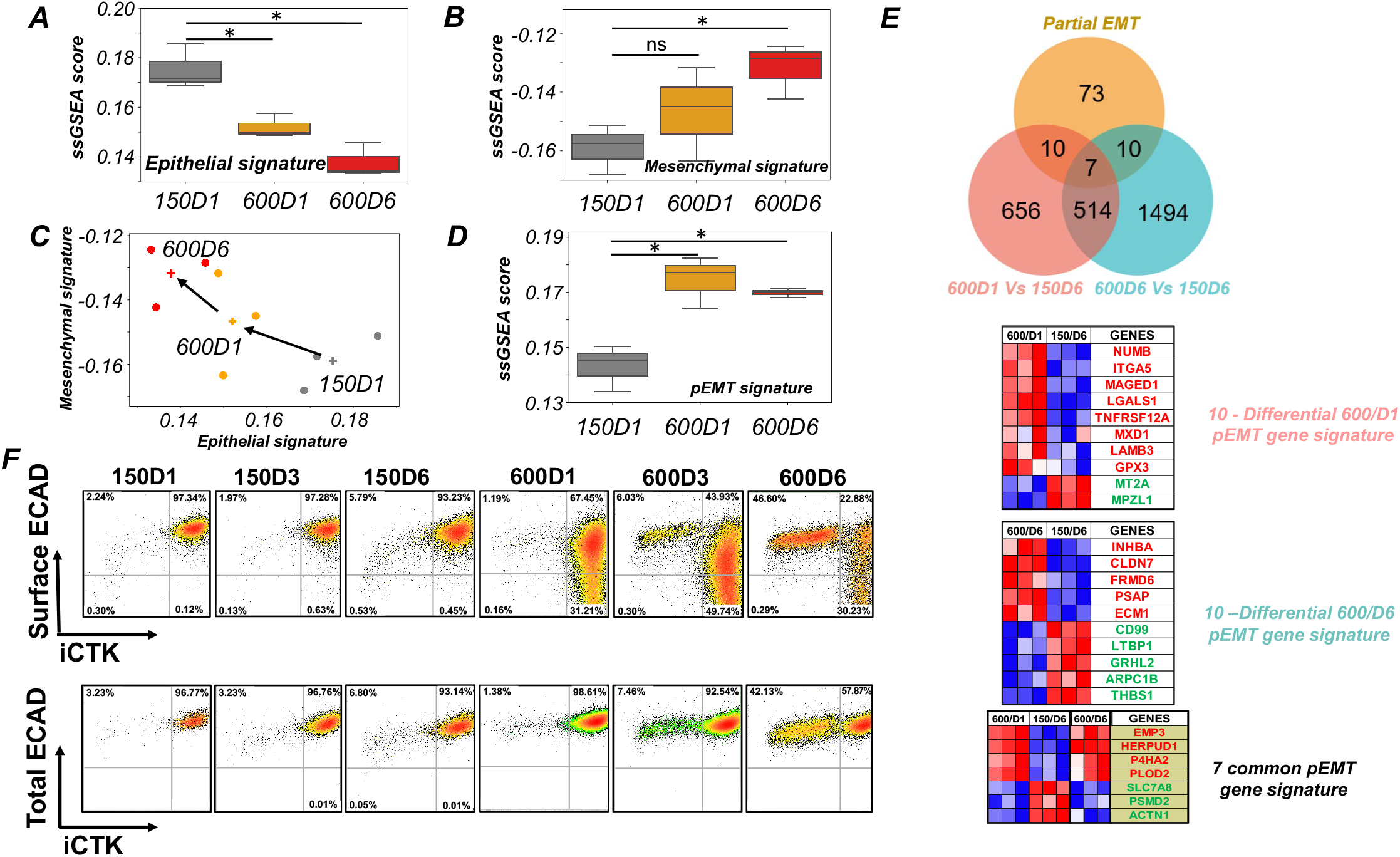
Large microtumors exhibit enrichment of partial EMT signature in comparison to small microtumors. **(A)** ssGSEA score analysis for ‘Epithelial tumor signature’ in 600/D1 and 600/D6 groups individually with 150/D6 microtumors as control shows significant decrease in epithelial expression over six-day period in both 600/D1 (*p*=0.01*) and 600/D6 (*p*=0.01*) groups; **(B)** ssGSEA score analysis for ‘Mesenchymal tumor signature’ in 600/D1 and 600/D6 groups individually with 150/D6 microtumors as control shows significant increase in expression of mesenchymal signatures over six day period in 600/D6 (*p*=0.01*) group but not in the 600/D1 group; **(C)** 2D EMT plots showing Epithelial-Mesenchymal characteristics of 150/D6, 600/D1 and 600/D6 microtumors. **(D)** ssGSEA score analysis for ‘partial EMT signature’ in 600/D1 and 600/D6 groups individually with 150/D6 microtumors as control shows significant increase in expression of partial EMT signatures [28] as early as day 1 in 600/D1 microtumors (*p*=0.01*) and is maintained over six-day period in 600/D6 (*p*=0.01*); **(E)** Venn diagram comparison of partial EMT signature set [28] with *600/D1 versus 150/D6* and *600/D6 versus 150/D6* group shows that total 17 genes are enriched in *600/D1 versus 150/D6* as well as in *600/D6 versus 150/D6* group with an overlap of 7 genes in both these groups. Heatmap shows the intensity values of genes from biological replicates in 600/D1, 600/D6 and 150/D6 group as separate list for 10 differentially regulated genes present only in 600/D1, 10 genes present only in 600/D6 and 7 genes common to both the groups. In the heatmap, ‘*red*’ shows upregulated status and ‘*blue*’ shows downregulated status. The corresponding gene names are illustrated with ‘*red*’ for upregulated status and ‘*green*’ for downregulated status; **(F)** Time-dependent expression of E-cadherin (E-CAD) and pan-cytokeratin (iCTK) in T47D cells in 150μm and 600μm microtumors over day 1, 3 and 6.

To identify partial EMT gene signatures in our breast microtumor models, we analyzed expression of partial EMT markers in 600/D1 and 600/D6 microtumors using available partial EMT gene signatures from head and neck cancer patients as a reference dataset [28]. We first calculated the enrichment of partial EMT signature using ssGSEA and found a significant increase in the partial EMT signature as early as 600/D1 compared to the 150/D6 tumors (**Figure 2D**). We then identified 17 partial EMT genes significantly enriched in 600μm microtumors as early as day 1and are referred to as ‘600/D1 partial EMT gene signature set’ (**Figure 2D**). Over the six-day period in culture, non-migrating 600/D1 microtumors transitioned to collectively migrating phenotype (600/D6). Through comparison of gene expression in 600/D6 versus 150/D6, we identified enrichment of 17 partial EMT genes, referred to as ‘600/D6 partial EMT gene signature set’ (**Figure 2D**). The Venn diagram of comparative analysis of ‘600D1 partial EMT gene signature set’ and ‘600D6 partial EMT gene signature set’ identified 10 genes differentially expressed only in 600/D1 and 10 genes differentially expressed only in 600/D6 (**Figure 2E**). Seven partial EMT genes were common to both 600D1 and 600D6 microtumors, of which *EMP3, HERPUD1, P4HA2, PLOD2* were upregulated and *SLC7A8, PSMD2, ACTN1* were downregulated in both 600/D1 and 600/D6 microtumors compared to control 150/D6 microtumors (**Figure 2E**). Of the 10 partial EMT genes differentially expressed only in 600D1 microtumors (**Figure 2E**), *NUMB, ITGA5, MAGED1, LGALS1, TNFRSF12A, MXD1, LAMB3*, and *GPX3* were most upregulated while MPZL1 and *MT2A* were downregulated (in descending order of expression) compared to 150/D6 microtumors. In ‘600/D6 partial EMT gene signature’, *INHBA* was the most upregulated gene followed by *CLDN7, FRMD6, PSAP* and ECM1 while *CD99, LTBP1, GRHL2, ARPC1B* and *THBS1* were top five downregulated genes (in descending order of expression) (**Figure 2E**).

Of note is the over-expression of some ‘phenotypic stability factors’ (PSFs) in 600/D1 microtumors and later in 600/D6 microtumors. The PSFs stabilize or maintain the partial EMT phenotype in tumor cells and prevent complete epithelial to mesenchymal transition (EMT) [45]. Recent studies have suggested the role of NUMB endocytic adaptor protein (*NUMB*), nuclear factor erythroid 2-related factor 2 (*NRF2*) and nuclear factor of activated T-cell (*NFATc*) as phenotypic stability factors [46, 47]. Our microarray expression profile of 600/D1 and 600/D6 microtumors revealed upregulation of *NUMB* [46] in hypoxic, but non-migratory 600/D1 microtumors supporting presence of partial EMT as early as day 1 (**Figure 2E**). In migratory microtumors (600/D6), *GRHL2* was found to be downregulated. While GRHL2 was initially proposed as a PSF [6], more recent studies have indicated that it is more of an MET inducer than a PSF [48–50], thus its downregulation during later time is expected, as cells lose their epithelial traits progressively (**Figure 2E**).

Observation that the partial EMT genes are expressed as early as day 1 and partial EMT phenotype is maintained over six-day period in large microtumors corroborates our previous work showing co-expression of E-CAD (epithelial marker) and VIM (mesenchymal marker) in migrating cells in large microtumors without the loss of E-CAD [21, 27]. To further understand the expression profile of E-CAD in our microtumor model, we evaluated surface and total (surface and intracellular) expression of E-CAD in small (150μm) and large (600μm) microtumors. In small non-migratory microtumors, the expression profile of E-CAD did not change from day 1 to day 3 to day 6 (**Figure 2F, top panel**). In large microtumors, we observed loss of surface E-CAD over six-day period, which is consistent with our prior work showing higher level of soluble E-CAD in the conditioned media of large microtumors [21]. However, analysis of combined surface and intracellular E-CAD revealed no change in the total E-CAD (**Figure 2F, bottom panel**). Interestingly, similar observations have been reported in pancreatic ductal adenocarcinoma (PDAC) model, highlighting the loss of epithelial phenotype through ECAD protein internalization instead of transcriptional repression of E-CAD, resulting in a partial EMT phenotype with cells showing gain of mesenchymal markers [51]. The study further reported that the cells expressing partial EMT migrate as clusters, which closely resembles the findings from our 3D breast microtumor model wherein large microtumors migrate as clusters and exhibit partial EMT phenotype. In a follow-up study using the same PDAC model system, calcium signaling – a key driver of the PSF, NFATc – was proposed to drive partial EMT [52], thereby strengthening our observations on some PSFs being differentially regulated as early as 600/D1.

### 3.3 Tumor-intrinsic hypoxia drives partial EMT and collective migration in large microtumors

To identify transcriptional signatures defining collective migration phenotype observed in the large hypoxic microtumors (600/D1, 600/D6) compared to small non-hypoxic microtumors (150/D6), we used the Gene Ontology gene signatures associated with tissue migration (GO: Tissue Migration) and evaluated the overlap between GO: Tissue Migration in 600/D1 *versus* 150/D6 and 600/D6 *versus* 150/D6 comparison groups (**Figure 3A**). Compared to 150/D6 microtumors, we identified 18 gene differentially expressed in non-migratory 600/D1 microtumors (denoted as ‘pre-migratory signature’) and 43 genes differentially expressed in collectively migrating 600/D6 microtumors (denoted as ‘migratory gene signature’). Of these differentially expressed genes, 11 genes were commonly expressed in both the groups, which maintained their expression over six-day period (**Figure 3A**). Of 18 genes, 7 genes were unique to non-migratory 600/D1 microtumors while out of 43 genes, 32 genes were unique to migratory 600/D1 microtumors (**Figure 3A**). We then calculated the ssGSEA scores for GO: Tissue Migration signatures and found significantly higher ssGSEA scores for the 600/D6 microtumors in comparison to 600/D1 microtumors (**Figure 3B**), suggesting that acquisition of migratory properties probably require acquisition of a late tumor-intrinsic hypoxia signatures as well as induction of the partial EMT program. This was further validated in the two-dimensional plots of Tissue Migration with Hallmark Hypoxia signature as well as with the partial EMT signatures (**Figure 3C and 3D**).

**Figure 3:**
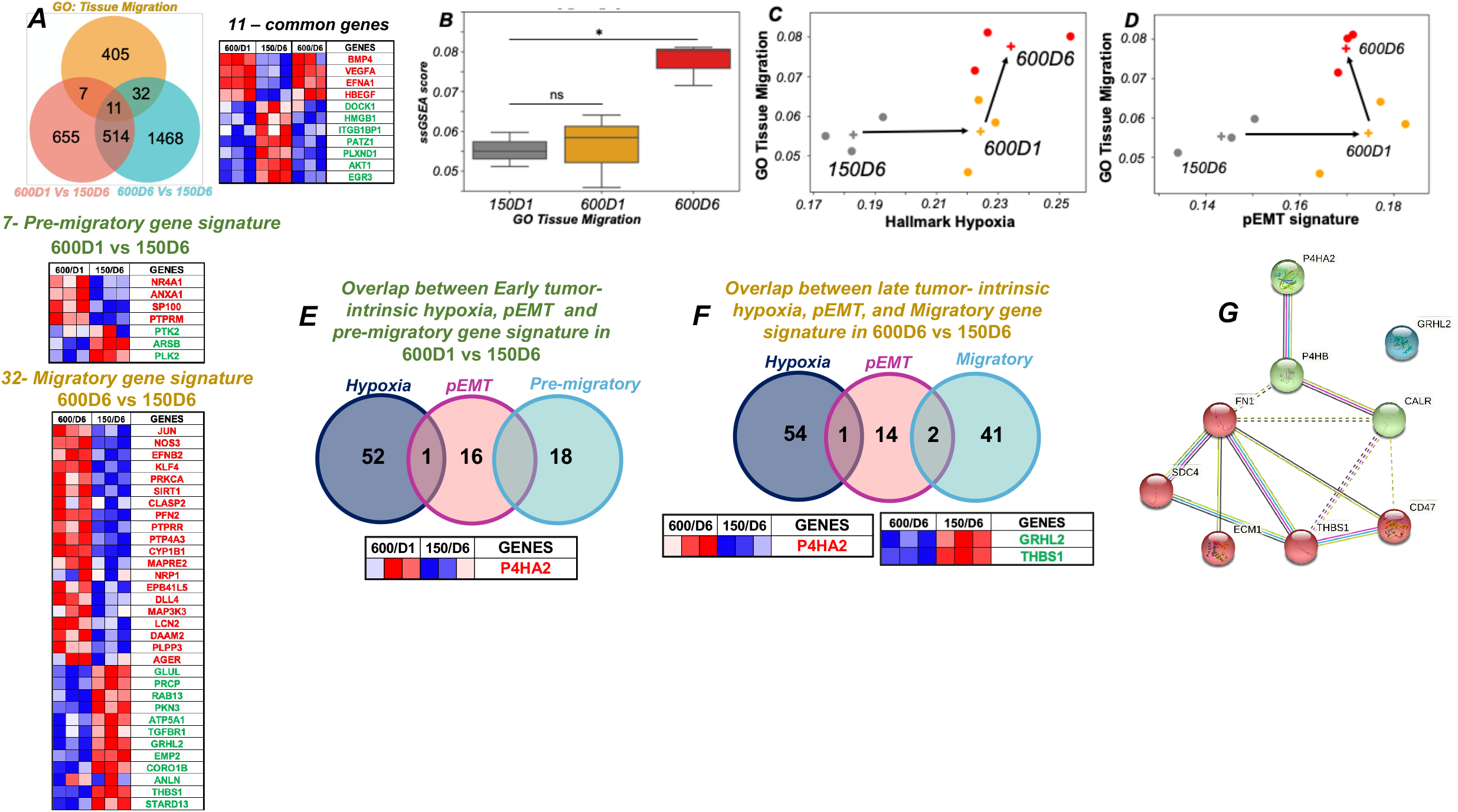
Tumor-intrinsic hypoxia drives partial EMT and collective migration in large microtumors. **(A)** Venn diagram comparison of *GO: Tissue migration* with *600/D1 versus 150D6* and *600D6 versus 150D6* comparison groups shows an overlap of 18 genes with *600/D1* (denoted as *pre-migratory gene signature*) and 43 genes with *600/D6 (denoted as migratory gene signature*). Eleven genes are common among *GO: Tissue migration*, pre-migratory and migratory gene signatures. Seven genes are differentially expressed only in *600/D1* and 32 genes are differentially expressed only in *600/D6*. These are represented as heat maps with corresponding intensity values of genes from biological replicates in 600/D1 and 600/D6 *versus* 150/D6 group. In the heatmaps, *‘red’* shows upregulated status and *‘blue’* shows downregulated status. The corresponding gene names are illustrated in ‘*red*’ for upregulated status and ‘*green*’ for downregulated status. **(B)** ssGSEA score analysis for *GO Tissue Migration* in 600/D1 and 600/D6 groups individually with 150/D6 microtumors as control shows significant increase in tissue migration only in 600/D6 (*p*=0.005*) group but not in the 600/D1 group; **(C)** 2D plot of *Hallmark Hypoxia* and *Tissue Migration* showing the trajectory of transition of the microtumors on the phenotypic space of hypoxia and tissue migration; **(D)** 2D plot of partial EMT phenotype and *Tissue Migration* showing the trajectory of transition of the microtumors on the phenotypic space of partial EMT and tissue migration; **(E)** Venn diagram comparison of early tumor-intrinsic hypoxia, partial EMT and pre-migratory gene signature set in in *600/D1 versus 150/D6* shows an overlap of one gene signature (*P4HA2*) between tumor-intrinsic hypoxia and partial EMT. Heatmap shows the intensity values of *P4HA2* from biological replicates in 600/D1 and 150/D6. The corresponding gene names are illustrated with *‘red’* for upregulated status **(F)** Venn diagram comparison of late tumor-intrinsic hypoxia, partial EMT and migratory gene signature set in *600/D6 versus 150/D6* shows an overlap of one gene signature (*P4HA2*) between tumor-intrinsic hypoxia and partial EMT, and two genes (*GRHL2, THBS1*) between partial EMT and migratory gene signature. Heatmap shows the intensity values of genes from biological replicates in 600/D6 and 150/D6 group. In the heatmap, ‘*red*’ shows upregulated status and *‘blue’* shows downregulated status. The corresponding gene names are illustrated with ‘*red*’ for upregulated status and *‘green’* for downregulated status. (G) Protein-protein interaction (PPI) network analysis of gene signatures common among hypoxia, partial EMT and tissue migration i.e. *P4HA2, GRHL2, THBS1*.

We further analyzed the overlap between hypoxia, partial EMT and tissue migration gene signatures in 600/D1 *versus* 150/D6 and 600/D6 *versus* 150/D6 groups to gain mechanistic insights into how tumor-intrinsic hypoxia contributes to driving partial EMT phenotype, which in turn, can modulate migratory phenotype. As shown in **Figure 3E**, only *P4HA2* gene was common between early tumor-intrinsic hypoxia and partial EMT gene signature in 600/D1 microtumors. Indeed, *P4HA2 is* known to be important for hypoxic adaptation and collagen biogenesis[53]. Not surprisingly, we did not find any overlap between partial EMT and pre-migratory gene signatures in 600/D1 microtumors (**Figure 3E**). This analysis supports our observations, wherein 600/D1 microtumors exhibit tumor-intrinsic hypoxia and partial EMT; however, they are yet to migrate. As large microtumors start migrating by day 3, we observed enrichment of both partial EMT and migration gene signatures in 600/D6 microtumors. This may suggest that tumor-intrinsic hypoxia is responsible for initiating a pre-migratory phenotype, but acquisition of partial EMT may be required to further induce collective migration. Indeed, further evaluation of the overlap between late tumor-intrinsic hypoxia and 600/D6 partial EMT gene signatures revealed *P4HA2* as a common gene shared between the two signatures (**Figure 3F)**. Thus, our analysis points to *P4HA2* as a central regulator of tumor-intrinsic hypoxia, partial EMT and migration. Few studies have highlighted role of *P4HA2* in driving EMT transition in glioma [54] and tumor migration in prostate cancer [55]; however, whether *P4HA2* drives migration through partial EMT states is not yet studied. Comprehensive gene expression analysis of hypoxic, yet non-migratory 600/D1 microtumors as well as that of hypoxic and migratory 600/D6 microtumors suggests that upregulation of *P4HA2* expression in response to tumor-intrinsic hypoxia plays a key role in inducing partial EMT states, which can further lead to collective migration.

We also identified two PSF genes signatures (*GRHL2* and *THBS1*) common between partial EMT and collective migration phenotype in 600/D6 microtumors (**Figure 3F)**. The role of *GRHL2*, another PSF found to be downregulated in 600/D6 microtumors, has been discussed above in Section 3.2*. THBS1* has been extensively studied to drive migration in cancer; however, its role as a partial EMT marker is not yet defined.

To further understand the protein-protein interaction (PPI) between these molecules, we performed *STRING* pathway analysis [56] (**Figure 3G**). No known interaction was found between *P4HA2* and *GRHL2*; however, the PPI network highlighted direct experimental regulation of P4HA2, which exhibit protein-protein interactions with migration related genes namely, *THBS1, FN1* and *ECM1*.

### 3.4 Inhibition of P4HA2 at early time point attenuates migration in large microtumors

Comparative analysis of common genes among early/late tumor-intrinsic hypoxia, partial EMT and migratory gene signatures (**Figure 3D-E**) revealed that hypoxia-responsive *P4HA2* gene was upregulated in 600//D1 (**Figure 3D**) and its expression was maintained over six-day period in 600//D6 (**Figure 3E**). Therefore, we hypothesized that early inhibition of *P4HA2* will attenuate collective migration in the large microtumors. To study the effect of *P4HA2* inhibition on collective migration, we treated large microtumors with 1,4-DPCA, a P4HA inhibitor starting from day 1 to day 6. The photomicrographs of representative microtumors **(Figure 4A)** illustrate comparative migratory phenotype in untreated (control) and 1,4-DPCA-treated 600μm microtumors in a time-dependent manner (D1, D3 and D6). In comparison to untreated microtumors, 1,4-DPCA-treated microtumors showed decrease in the migratory front or the distance of migration travelled by the microtumors from the edge of the well. The quantitative analysis of migrating microtumors post-1,4-DPCA treatment showed reduction in the number of migratory microtumors as early as day 3 (*p*=0.0084**), which was sustained over 6 days (*p*=0.011*) when compared to untreated microtumors (**Figure 4B**). We also measured the distance of migration in the migratory microtumors treated with 1,4-DPCA in comparison to controls. The distance of migration was significantly reduced in 1,4-DPCA-treated microtumors compared to untreated controls on day 3 (*p*=0.001***) and day 6 (*p*=0.003**) (**Figure 4C**). Quantitative RT-PCR studies showed significant decrease in the expression of both *P4HA1* and *P4HA2* in 1,4-DPCA treated microtumors compared to controls (**Figure 4D**), indicating that 1,4-DPCA is a P4HA inhibitor that is not specific for *P4HA2*. Together, results from inhibition studies strongly suggest P4HA inhibition as an effective treatment strategy to block collective migration.

**Figure 4:**
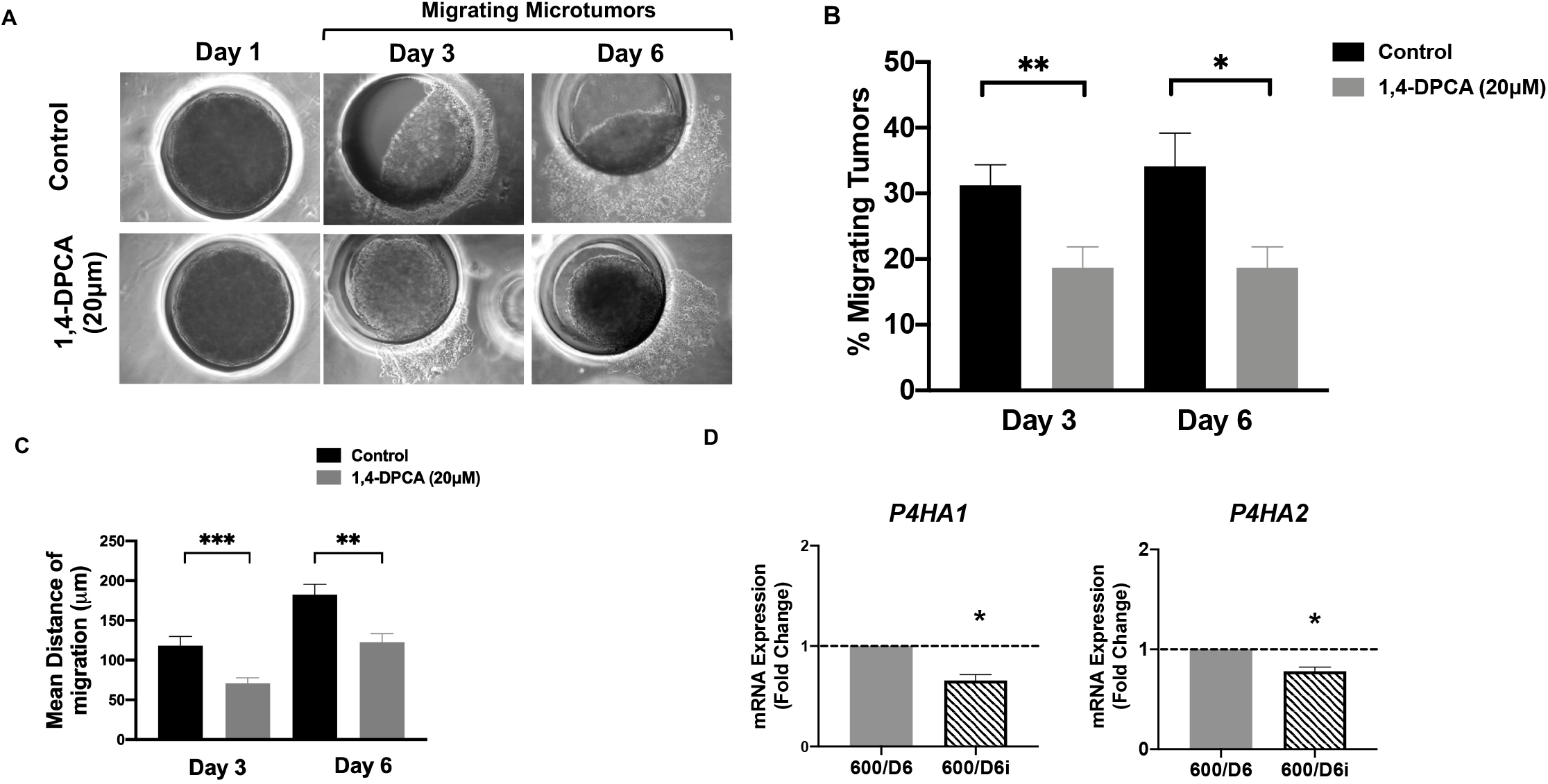
Inhibition of P4HA2 at early time point attenuates migration in large microtumors. **(A)** Photomicrographs of migrating 600μm microtumors in untreated (top panel) and 1,4-DPCA-treated (bottom panel) devices over day 1, day 3 and day 6; **(B)** Percentage migrating microtumors on 600/D3 and 600/D6 in untreated and 1,4-DPCA treated 600μm microtumors; **(C)** Effect of 1,4-DPCA on distance of migration travelled by 600μm microtumors evaluated on day 3 and day 6 for 10 individual tumors. 1,4-DPCA significantly reduced distance of migration on day 3 and day 6 as compared to untreated microtumors; **(D)** Effect of 1,4-DPCA inhibition on common hypoxia and pEMT signature (P4HA2) in 600μm microtumors w.r.t 600/D6. Fold change (mRNA expression) for P4HA1 and P4HA2 was determined by quantitative real-time PCR for 600μm microtumors (D6) harvested on day 6 in untreated and 1,4-DPCA treated 600μm microtumors. GAPDH, the housekeeping gene was used as a reference gene. Relative expression of P4HA1 and P4HA2 was compared to untreated 600/D6 controls (dotted line parallel to x-axis on bar graph). P4HA1 and P4HA2 expression decreased significantly in 1,4-DPCA treated 600μm microtumors. Data is represented as mean ± SEM (Unpaired t-test, * p < 0.05, ** p < 0.01, *** p < 0.001 w.r.t. untreated 600/D6).10

## 4. Discussion

With updated guidelines in the field of EMT, a current focus is to understand the transiently stable partial EMT phenotypes wherein malignant cells acquire mesenchymal characteristics without a complete loss of epithelial markers [2]. Role of tumor microenvironmental factors such as tumor-intrinsic hypoxia, extracellular matrix or stromal factors in inducing or maintaining the partial EMT state remains an important understudied area of research [1]. Tumor-intrinsic hypoxia is one of the key contributors, if not the key driver and has long been known to drive tumor invasion and migration.

We have developed unique microtumor models that recapitulate tumor-intrinsic hypoxia using control over microtumor size. Small 150μm microtumors remain non-hypoxic and non-migratory over six-day culture period while large 600μm microtumors develop hypoxic signatures as early as day 1 [27] and exhibit collective migration starting from day 3 in culture [21]. We have also shown that once acquired, the migratory phenotype is irreversible [20] and that HIF-1 inhibition at later time (after day 3) cannot block collective migration [21]. In this report, we exploited these unique microtumor models to investigate the link between tumor-intrinsic hypoxia, partial EMT and collective migration using integrated computational and experimental approaches.

From gene expression analysis, we propose a unique 53 gene-early tumor-intrinsic hypoxia gene signature and 55 gene-late tumor-intrinsic hypoxia gene signature. To the best of our knowledge, this is one of the first attempts to identify transcriptional changes in real time in response to naturally induced tumor-intrinsic hypoxia at different time points as 3D microtumors transition from non-migratory to migratory phenotypes. We also identify transcriptional markers for partial EMT phenotype and propose a unique 17 gene-partial EMT gene signature for non-migratory 600/D1 and 17 gene-partial EMT gene signature for migratory 600/D6 microtumors. The ssGSEA analysis highlights phenotypic plasticity of T47D cells, which exhibit predominantly epithelial phenotype when cultured as non-hypoxic 150μm microtumors, epithelial-like partial EMT phenotype in hypoxic 600μm microtumors (day 1) transitioning to more mesenchymal-like partial EMT phenotype on day 6 as hypoxic 600μm microtumors become migratory. The analysis of overlap between tumor-intrinsic hypoxia, partial EMT and migration signatures at early and late time-points suggests *P4HA2* as a key molecule, which is expressed early in non-migratory hypoxic microtumors and is enriched further as non-migratory microtumors transition into migratory phenotypes. Thus, we identify *P4HA2* as a partial EMT marker induced by tumor-intrinsic hypoxia, and demonstrate its role in collective migration through pharmacological inhibition studies.

Our findings are supported from few other studies in literature. For example, in glioma cells, *P4HA2* promoted EMT via collagen-dependent PI3K/AKT pathway and *P4HA2* knockdown inhibited migration, invasion and EMT [54]. Collagen-stroma interactions in the ECM were shown to induce P4HA2 expression via HIF-1α [57]. In breast cancer patient microarray data, high mRNA expression of *P4HA2* was associated with collagen deposition (*COL1A1, COL3A1* and *COL4A1*) and poor patient outcomes [58]. Similar findings were reported in DCIS patients (pure *versus* mixed and invasive carcinoma) and *P4HA2* was suggested as an independent predictor of DCIS risk stratification [59].

## 5. Conclusion

Taken together, using our unique 3D microtumor models, we show that tumor-intrinsic hypoxia changes transcriptional landscape driving partial EMT and collective migration. We identify *P4HA2*, known to be important for hypoxic adaptation and collagen biogenesis, as a key regulator of tumor-intrinsic hypoxia, partial EMT and collective migration, which may also serve as potential target to inhibit acquisition of migratory phenotype in breast cancer cells.

